# Interference at Influenza Hemagglutinin Antigenic-Sites Determines Antibody Levels and Specificities after Repeat Vaccination

**DOI:** 10.1101/2021.06.01.446556

**Authors:** Christopher S. Anderson, David J. Topham

**Affiliations:** Department of Pediatrics, University of Rochester Medical Center.; David Smith Center for Vaccine Biology and Immunology, Department of Microbiology and Immunology, University of Rochester School of Medicine and Dentistry, Rochester, NY.

**Keywords:** influenza, vaccines, immune focusing, Antigenic Distance Hypothesis, Hemagglutinin, Stalk, pH1N1, Shape Space

## Abstract

**Background:** It is recommended that children receive a dose of the influenza vaccine at 6 months of age and a second dose the following season. In some years, the second dose will be the same vaccine formulation, in others years it with be re-formulated to include HA proteins derived from antigenically drifted or shifted circulating influenza strains. In addition, natural exposure to influenza can create permanent changes to the memory B cell repertoire and specificities. The effect that the specificity of pre-existing humoral immunity has on antibody levels and specificity after repeat vaccination is an ongoing research area.

**Methods:** We used a computational framework (ssMod.v1) to simulate scenarios that occurred during the 2009 influenza pandemic: children receiving a second dose who have previously exposed to the 2009 influenza HA antigen, children previously exposed to an HA antigen antigenically similar to the 2009 influenza HA antigen, and children previously exposed to an antigenically dissimilar strain. To assess the contribution of pre-existing immunity, two experimental permutations in the sMod.v1 were made: elimination of antibody-mediated antibody clearance of vaccine (HA) antigen by pre-existing antibodies and elimination of the ability of memory B cells to form germinal centers. In the simulation, 30 days after repeat vaccination antibody specificities were examined against 12 antigenically historical vaccine/HA variant influenza strains for each of the five canonical antigenic-sites and the subdominant, conserved, stalk antigenic-site.

**Results:** We found that elimination of antibody-mediated antigen clearance significantly increased antibody levels while elimination of the ability of memory B cells to form germinal center reactions significantly decreased the total antibody levels, but this was dependent on the antigenic relationship between the original vaccine and repeat vaccine and the particular antigenic-site. Moreover, highly-cross-reactive antibody was highest when the antigenic distance between original vaccine HA antigen and repeat vaccine HA antigen was larger and antibody-mediated antigen clearance was eliminated.

## Introduction

Each year in the United States, influenza virus infection leads to hospitalization in about 1 in every 1000 children, typically in those preschool and younger[1,2]. Influenza-specific antibodies are essential for protection against influenza infection[3]. In the United States, the Influenza virus vaccine is recommended for all children starting at six months of age in order to induce antibodies against the influenza surface protein, hemagglutinin (HA), which is highly correlated with protection against severe influenza disease[4]. Antigenic drift and antigenic shift of the HA protein in circulating viruses reduces the immunity afforded by the vaccine. Vaccine induced antibody has been shown to decrease over time and children who get influenza infection have lower antibodies specific to the circulating strain HA [5]. Therefore, the influenza vaccine is recommended each year after the original immunization. Some years the influenza vaccine will contain identical HA proteins to the previous year’s vaccine and some years it will have a new formulation with HA proteins derived from newly circulating antigenically drifted or shifted influenza strains.

Repeat vaccination studies have shown that prior influenza vaccination can reduce the effectiveness of the vaccine[6–10]. Multiple studies have demonstrated that this reduced vaccine effectiveness depends on the antigenic relationship between the original immunizing influenza vaccine HA protein and the revaccination HA protein results in differences in antibody specificities, leading to variation in vulnerability to circulating influenza viruses[11–16]. Individuals tend to have the highest level of circulating antibodies and memory B cells specific to influenza virus strains that circulated when they were young compared to more recent strains [9,17]. Upon repeat vaccination, it has been shown that the antibody and memory B cell specificity differs between individuals with different antibody specificities prior to repeat vaccination [5,9,10].

In 2009, a world-wide pandemic occurred due to a antigenic shift in the influenza virus in swine and subsequent transmission to humans[18]. Within a year a new influenza vaccine was formulated with proteins derived from the shifted strain and administered to the public, including young children. Research into antibody induced by the 2009 pandemic virus vaccine showed induction of highly-cross-reactive antibodies able to bind a broad range of antigenically distinct seasonal viruses and highly divergent avian influenza strains. Interestingly, the boost in cross-reactive antibody was only seen in younger individuals [19,20]. Furthermore, the increase in cross-reactivity in young individuals was found to be associated with the antigenic-sites to which antibodies were induced[20,21], with young individuals inducing antibodies towards conserved regions on the HA protein, including the stalk region, while older individuals had greater antibody specific for the 2009 H1N1 HA protein [20,22].

Many immunological agents have been implicated as the root of the dissimilarities in antibody specificities seen after repeat vaccination including regulatory T cells, memory B cells, and antibody. Immune processes have also been implicated such as the masking of novel epitopes by antibody, Fcγ receptor-mediated inhibition of B cell activation, and antibody-mediated clearance of antigen[5,6,8,12,23,24]. Early computational models suggested that both memory B cells and antibodies affect the antibody specificity of the humoral responses after repeat vaccination in a manner dependent on the antigenic distance between the influenza HA proteins included in the original vaccine and repeat vaccine[12]. Detailed computational models of the antibody-antigen interaction demonstrated that pre-existing antibodies play a crucial role in the specificities of antibody induced after repeat vaccination[25]. Pre-existing memory B cells have also been implicated in focusing the antibody response to conserved regions of the HA protein, in a manner dependent on the specificity of the pre-existing memory B cells[13,14]. We hypothesize that the specificity of pre-existing antibodies and memory B cells is a major determinant of the magnitude and specificity of the response to subsequent vaccination.

Here we test the effect of pre-existing memory B cells and antibodies, and their specificities on antibody responses to the 2009 H1N1 pandemic vaccine. Using the ssMod.v1 computational framework, which simulates humoral immune responses to historical/vaccine strain influenza HA protein antigens[26], we investigated the role pre-existing memory B cells and antibodies play in the specificity of the antibody response induced in children receiving their second dose of the influenza vaccine.

## Results

Three scenarios were simulated using the ssMod.v1 computational framework: (Scenario 1) children originally vaccinated with the 2009 H1N1 pandemic virus vaccine, (Scenario 2) children originally vaccinated with a formulation containing HA protein antigens antigenically similar, but not identical, to the 2009-pandemic virus HA protein in the vaccine, (Scenario 3) children originally exposed to a HA protein highly antigenically dissimilar to the 2009 H1N1 vaccine HA protein antigen. One “year” after the original vaccination a second vaccine dose, which was re-formulated to include the HA protein derived from the 2009 H1N1 pandemic virus, was administered (Figure 1).

**Figure 1.**
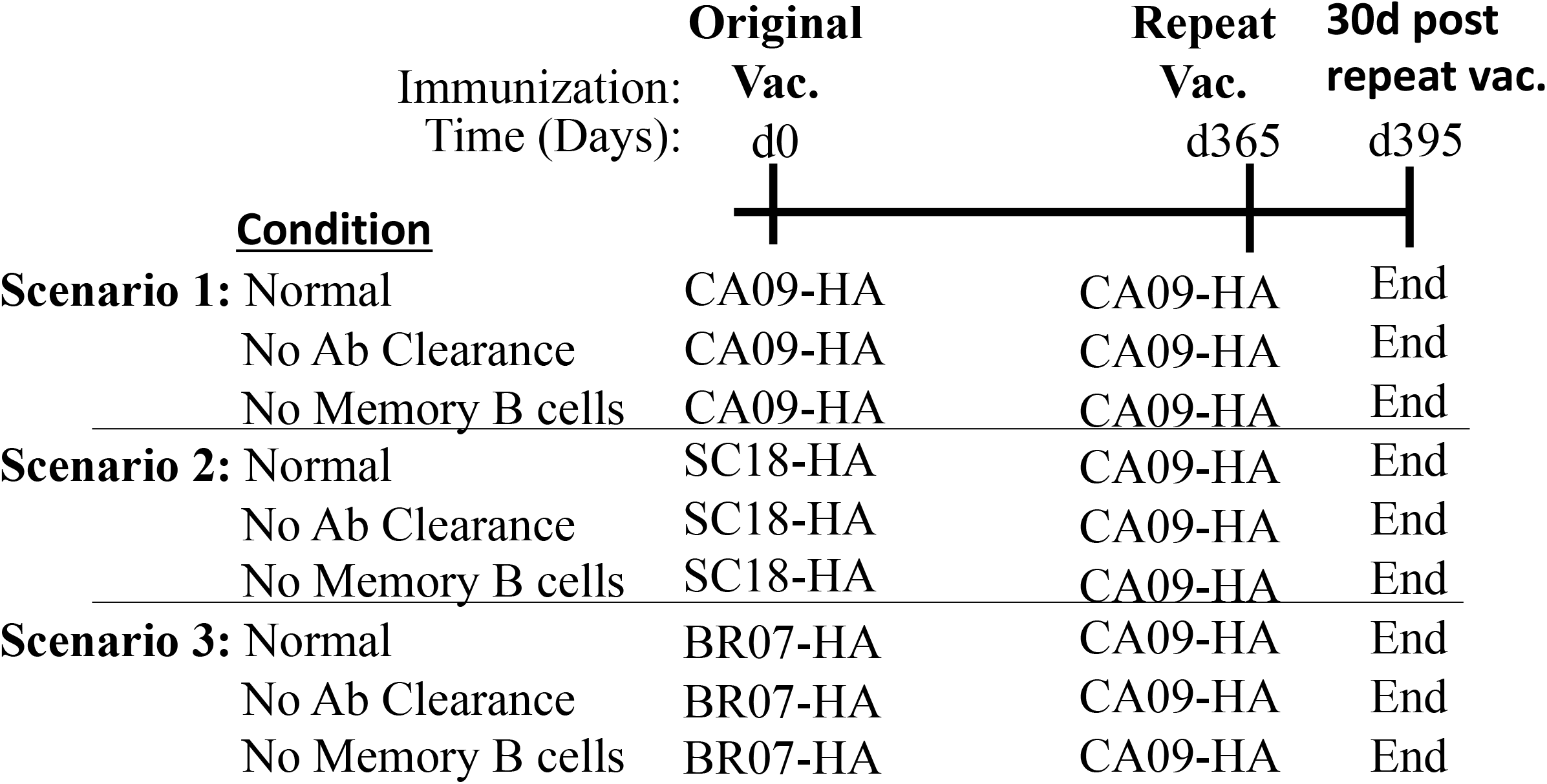
Schematic of Repeat Vaccine Scenarios and Perturbations. Three types of computational simulations were performed using the ssMod.v1 framework, Scenario 1-3. In each scenario a different HA antigen protein was given and one year later the 2009 pandemic HA antigen was given. For each scenario, two perturbations to the ssMod.v1 framework were made either removing the ability of antibody to clear antibody (No Ab Clearance) or removing the ability of memory B cells to become stimulated and form germinal centers (No Memory B cells).

Both pre-existing memory B cells and antibody have been shown to interfer with secondary immune responses. Therefore, for each scenario modeled, two perturbations were made in the simulations: the ability of antibodies to clear vaccine HA protein antigen was eliminated, resulting in antigen only diminishing from natural decay, and elimination of the ability of memory B cells to become activated and form germinal centers, resulting in only naive B cells contributing to germinal centers and thus producing antibody secreting cells.

The computational framework was sensitive to both pertubations.Total antibody levels 30 days post repeat vaccination were similarly affected by perturbations. In all scenarios, removal of antigen clearance by antibody significantly increased the total antibody levels compared to the unperturbed, normal, simulations. Removal of memory B cells from germinal center reactions significantly decreased the total antibody levels compared to unperturbed simulations for all three scenarios (Figure 2). Taken together, pre-existing antibody negatively interfered with total antibody responses while pre-existing memory B cells positively interfered with total antibody levels after repeat vaccination regardless of the antigenicity of the original exposure HA protein antigen.

**Figure 2.**
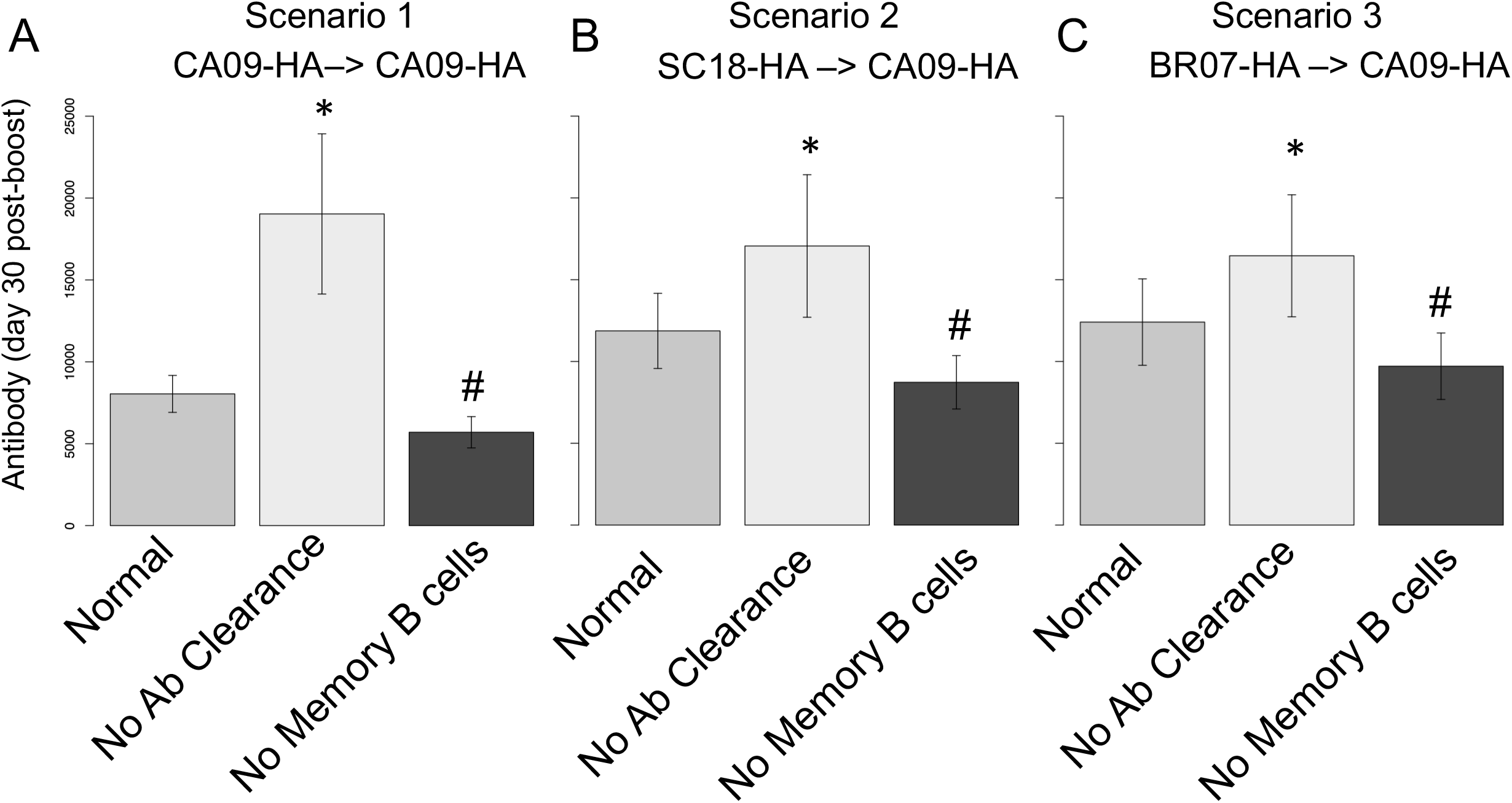
Total Antibody Levels Post Repeat Vaccination. Total antibody levels 30 days after repeat exposure to CA09-HA for simulations originally exposed to (A) CA09-HA, (B) SC18-HA, or (C) BR07-HA. The average of 50 simulations for each scenario for unperturbed (Normal) and perturbed (No Ab Clearance, No Memory B cells) conditions were calculated and error bars represent standard deviation. ‘*’ represents p-value less than 0.05 for two-sample t-test between Normal and No Ab Clearance antibody levels. ‘#’ represents p-value less than 0.05 for two-sample t-test between Normal and Memory B cell levels.

Elimination of antibody clearance affected cross-reactivity of antibodies induced 30 days after repeat vaccination. Cross-reactivity was determined against a wide-range of antigenically distinct, prototypical, HA antigens representing historical and seasonal-vaccine influenza strains. Removing antibody-mediated antigen clearance increased antibody cross-reactivity after repeat vaccination to all HA antigens, for both Scenario 1 and Scenario 3. When the repeat vaccine was antigenically similar, but not identical, to the first vaccine HA protein antigen removal of antibody-mediated antigen clearance only affected antibody binding for strains antigenically similar (Figure 3).

**Figure 3.**
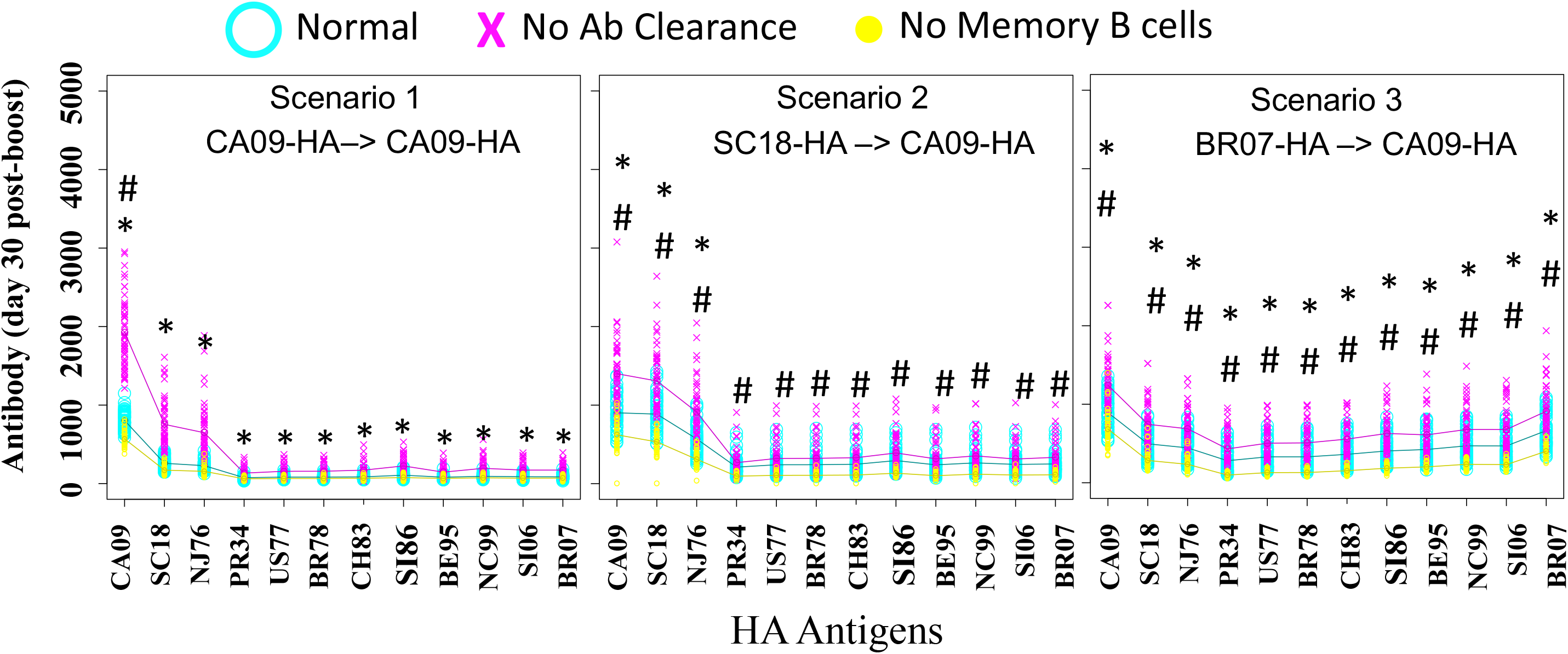
Strain Specific Antibody Levels After Repeat Vaccination. Antibody levels 30 days after repeat vaccination to HA antigens representing 12 antigenically distinct historical/vaccine influenza strains for each scenario. Antibody for normal, unperturbed, simulations are in blue open circle, No Ab Clearance are in red ‘X’, and No Memory B cell simulation antibody levels are marked by yellow full circles for each of the 50 simulations performed for each scenario and condition. ‘*’ represents p-value less than 0.05 for ANOVA between Normal and No Ab Clearance antibody levels. ‘#’ represents p-value less than 0.05 for ANOVA between Normal and Memory B cell levels.

Removal of the memory B cell’s ability to form germinal centers reduced antibody levels to all HA protein antigens (Figure 3), but only when a heterologous HA protein antigens were used (Scenario 2&3). Alternatively, when homologous antigens were used (Scenario 1), removal of memory B cell’s ability to form germinal centers had no significant effect on the cross-reactivity. Taken together, antibody-mediated clearance of antigen significantly altered cross-reactivity for all scenarios, while memory B cells only contributed to cross-reactivity of the antibody response when the repeat HA protein was antigenically distinct from the original HA protein antigen.

The number of highly-cross-reactive antibodies, those able to bind 10-12 HA protein antigens, was determined. Elimination of antibody-mediated antigen clearance for both Scenarios 1 & 3 significantly increased antibody levels, but not for the Scenario 2, which was not significantly affected. Eliminating memory B cells from germinal center reactions resulted in a significantly decreased the number of highly-cross-reactive antibodies induced in both heterologous HA protein antigen re-exposure scenarios (Scenario 2 & 3), but not for the homologous HA protein antigen re-exposure scenario (CA09-HA->CA09-HA). Taken together, the effect of pre-existing antibodies and memory B cells had on a number of antigenically distinct antigens each antibody was capable of binding after secondary exposure to HA antigen was largely dependent on the original exposure HA protein antigen.

The number of antibodies specific to each of the six HA antigenic-sites was determined. Repeat vaccination with homologous HA protein antigen (Scenario 1) was significantly affected by antibody-mediated antigen clearance, with all antigenic-sites showing a significant increase in antibodies level, including the subdominant Stk antigenic-site compared to unperturbed (Figure 5). For Scenario 2, elimination of antibody-mediated antigen clearance significantly increased antibodies levels to the three most conserved HA head antigenic-sites, but not to head antigenic-sites with larger antigenic distances or the subdominant, fully-conserved, Stk antigenic-site. Elimination of antibody clearance for the Scenario 3 simulations significantly increased antibody levels to antigenic-sites with very large (>7) antigenic distances and to the Stk antigenic-site, but not to the most conserved head antigenic-site, Ca1. Taken together, elimination of antibody-mediated antigen clearance increased antibodies to all antigenic-sites when the antigenic-distance between the original HA antigen and the repeat HA antigen was short, but when the antigenic-distance was increased, the effect of pre-existing antibodies depended on both the conservation and immunodominance of the antigenic-site.

**Figure 4.**
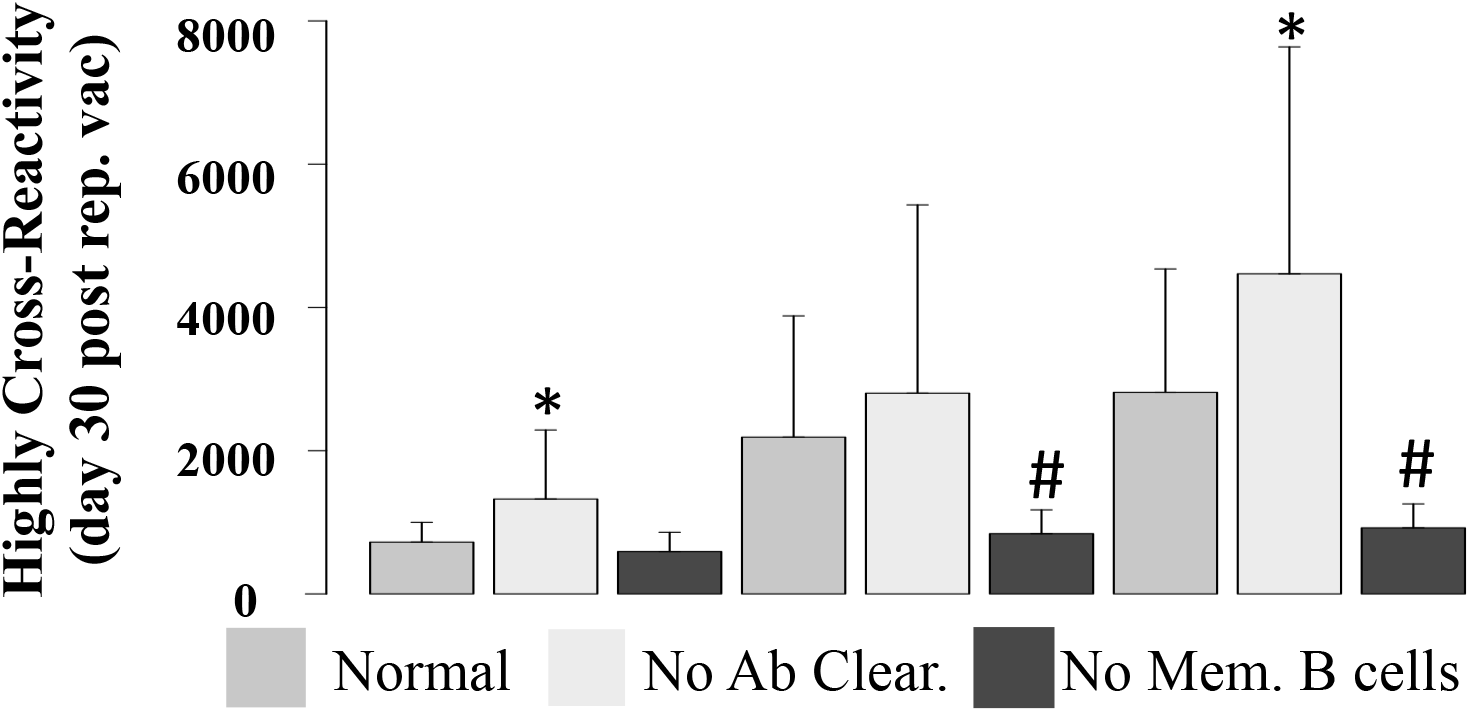
Cross-reactivity of Individual Antibodies. The number of HA protein antigens in which an antibody had affinity for most HA antigens(10- 12 HA protein antigens) was determined for each antibody present at day 30 post repeat vaccination. Barplots represents the average of 50 simulations performed for each scenario and condition. Error bar represents standard deviation. ‘*’ represents p-value less than 0.05 for two-sample t-test between Normal and No Ab Clearance antibody levels. ‘#’ represents p-value less than 0.05 for two-sample t-test between Normal and Memory B cell levels.

**Figure 5.**
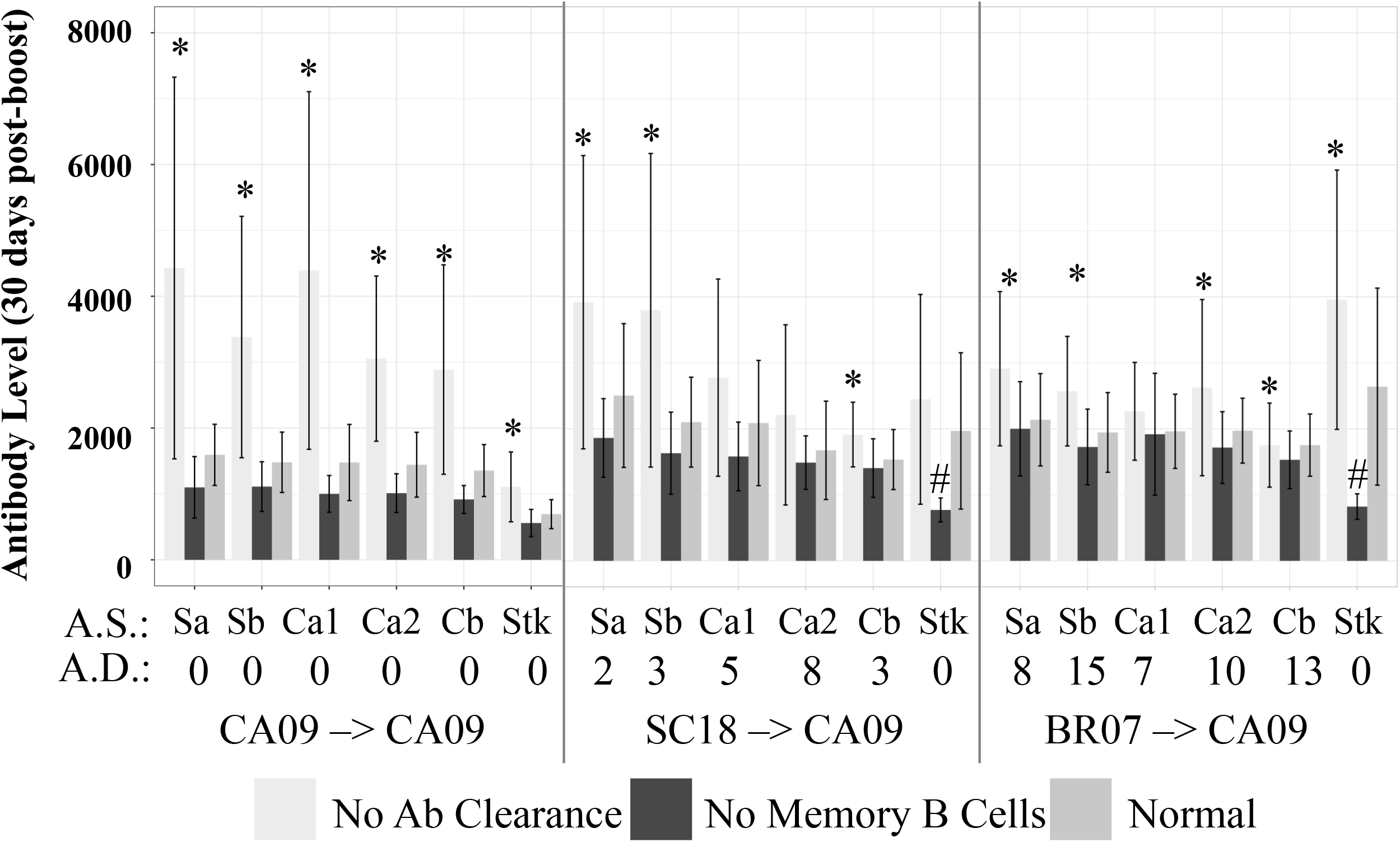
Antigenic-Site Specific Antibody Levels. The number of antibodies able to bind the five canonical head antigenic-sites and a conserved (Stk) site was determined at day 30 post repeat vaccination. Barplots represents the average of 50 simulations performed for each scenario and condition. Error bar represents standard deviation. ‘*’ represents p-value less than 0.05 for ANOVA between Normal and No Ab Clearance antibody levels. ‘#’ represents p-value less than 0.05 for ANOVA between Normal and Memory B cell levels. The antigenic-distances (0-20 A.D.) between original HA protein antigen and repeat HA protein antigen for each antigenic-site (A.S.) is shown under each barplot.

Elimination of memory B cell’s ability to form germinal centers had no significant effect on antibody levels to head antigenic-sites 30 days after repeat exposure to HA protein antigen regardless of the antigenic distance between the HA protein antigens (Figure 5). Alternatively, removal of memory B cells from the germinal center reactions significantly decreased antibody to the Stk antigenic-site to both scenarios with re-exposure to heterologous HA antigen. Taken together, memory B cells played a significant role during repeat vaccination, but only during heterologous HA antigen re-exposure and only in the conserved, subdominant, antigenic-site.

## Methods

### Determination of antigenic distance between HA antigens

As previously described[27], HA amino acid sequence data was obtained from the Influenza Resource Database (fludb.org) [28]. In short, a viral-sequence-based antigenic distance calculation is used to determine the antigenic distance between canonical antigenic-sites of the HA antigen. Virtual HA antigens in the framework were then defined in a manner that captured the antigenic relationship between antigens. In additional to the five canonical antigenic sites (Sa, Sb, Ca1, Ca2, Cb), a single, fully-conserved, stalk antigenic site (Stk) was also represented in the framework. Twelve HA antigens were represented in the model using HA genome sequences from the Influenza Resource Database (fludb.org) [28]: A/California/07/2009 (CA09) [NC_026433], A/Brisbane/59/2007 (BR07) [KP458398], A/South Carolina/01/1918 (SC18) [AF117241], A/Beijing/262/1995 (BE95) [AAP34323], A/Brazil/11/1978 (BR78) [A4GBX7], A/Chile/1/1983 (CH83) [A4GCH5], A/New Caledonia/20/99 (NC99) [AY289929], A/Singapore/6/1986 (SI86) [ABO38395], A/Solomon Islands/3/2006 (SI06) [ABU99109], A/USSR/90/1977 (US77) [P03453], A/New Jersey/11/1976 (NJ76) [ACU80014], A/Puerto Rico/8/1934 (PR34) [HQ008261].

### Computational Framework

Antigenic distances were used to define antigens in the open-source computational framework, ssMod.v1[26]. ssMod.v1 is an agent-based modeling system which explicitly models the antigenic-sites of virus proteins and the immunoglobulin-binding-domains of B cells and antibodies in an immunological shape space [12,29,30]. The framework captures the distributions of the paragenic properties of the B cell receptor binding region using a Lazy Evaluation approach[31].

### Simulating Repeat Vaccination Scenarios

Three scenarios were simulated using the ssMod.v1 computational framework. In Scenario 1, ssMod.v1 framework was set to simulate vaccination with HA protein antigen derived from the 2009 pandemic strain (CA09-HA) and repeat HA protein antigen exposure to the same (homologous) antigen one year later. In Scenario 2, ssMod.v1 was set to simulate vaccination with HA protein from the 1918 pandemic strain HA protein antigen (SC18-HA) and repeat HA protein antigen exposure with the moderately antigenically similar 2009 pandemic HA protein antigen. In Scenario 3, ssMod.v1 was set to simulate vaccination with the 2007 vaccine strain HA antigen (BR07-HA) and repeat exposure with the highly antigenically distinct 2009 pandemic HA protein antigen. Antibody numbers and specify was tracked throughout the simulation.

## Discussion

In 1977, the seminal work by Perelson et al. described how the binding domain (paratope) of an immunoglobulin receptor can be thought to exist in a multi-dimensional space where the position of the paratope in that space is defined by the biochemical properties (e.g. hydrogen bonding, Van Der-waals, hydrophobicity) that determine the ‘shape’ of the paratope. Paratopes with similar shapes are closer together in the ‘shape space’ while dissimilar ‘shapes’ are further apart in the shape space. Epitopes, regions of a protein bound by a paratope, also exist in ‘shape space’ such that an epitope with a high affinity for the paratope can be thought to reside close to that paratope in the ‘shape space’. Perelson et al. also described the ‘ball of stimulation’ of a paratope since each paratope can bind epitopes over a range of affinities and therefore can bind a slightly antigenically-drifted epitopes. Here we used a computational framework (ssMod.v1) based on the principals of shape space.

In 1999, Smith et al. derived the parameters for the immunological shape space in order to understand how repeat vaccination can reduce vaccine efficacy when comparing to first-time vaccine receivers, and why this only occured during some influenza epidemics. Smith et al. results suggested that the creation of antibody and memory B cells during initial exposure resulted in increased numbers of paratopes (antibodies and memory B cells) in certain areas of the shape space resulting in negative interference, where antibodies and memory B cells interfere with boosting of antibody responses, resulting in a decrease in the expansion of new paratopes during repeat vaccination, potentially leading to increased susceptibility to a drifted influenza virus. The increase in paratopes also results in positive interference where pre-existing cross-reactive antibody and pre-existing memory B cells produced in the response to the original vaccine HA antigen can boost the antibody levels to slightly-drift HA protein antigens. Known as the Antigenic Distance Hypothesis, Smith et al. showed that reduced efficacy during repeat vaccination was dependent on where in the shape space the epidemic strain expands paratopes, with outcomes being worse if the distance between the two vaccine HA antigens was close in shape space, but the epidemic strain is far, but not when the two vaccine HA antigens were very different. Our results support and expand these results demonstrating that positive and negative interference occurs during repeat exposure, but can be different for different antigenic-sites and results in differences in cross-reactivity. For instance, we showed that if positive interference occurred at a conserved antigenic site an increase in cross-reactive antibodies occurs.

Real-world studies have supported the Antigenic Distance Hypothesis and our current findings. Originally studies describing ‘Original Antigenic Sin’ demonstrated using a set of antigenically drifted influenza strains that the highest antibody levels in an individual were against strains that circulated in the first few years of a person’s life. This suggests that the ability of an individual’s shape space to expand antibody paratopes is greater during initial exposure to the virus than subsequent exposures. Fonville et al. used an approached known as antigenic cartography to defined a composite shape space where each HA antigen, but not epitope, are represented in a space. Using this approach, Fonville et al. overlayed an individual’s antibody binding data for each HA antigen represented in the shape space and demonstrated signatures of original antigenic sin prior to infection, with negative interference occurring when the virus strain was well matched to existing antibody, consistent with our results.

Pre-existing memory B cells have also been shown to demonstrate similar signatures of original antigenic sin[9] and multiple studies have suggested that when the antigenic distance between original and repeat HA antigens is larger, memory B cells complementary to the regions conserved between the HA antigens preferentially expand resulting in expansion of paratopes (antibodies) complementary to these conserved regions, sometimes referred to as ‘antibody focusing’[13,14]. Here we showed that pre-existing antibodies and memory B cells both can affect antibody levels after vaccination. The effect occurred in a manner dependent on the conservation of HA antigenic-site (epitopes) and the individual’s expansion of certain memory B cells and antibodies (paratopes) specific to certain HA antigenic-sites (epitopes) in shape space during original HA protein antigen exposure. The presence of pre-existing, cross-reactive, antibody generally reduced the antibody response to HA antigenic-sites, but not always, and depended on both immunodominance of the antigenic-site and its antigenic distance. Pre-existing antibody during homologous vaccination had a greater impact than antibody during heterologous vaccination with drifted strains, consistent with Smith et al. They also support studies showing increased antibody levels to subdominant epitopes when the antigenic distance between the original vaccine and repeat vaccine HA antigens is large. Moreover, in our simulations, the elimination of antibody-mediated antigen clearance significantly increased total antibody levels. These findings are consistent with recent studies by Zarnitsyna et al. which suggest that masking of epitopes by pre-existing antibodies, decreases B cell activation, resulting in a decrease in cross-reactivity[25]. Furthermore, our results suggest that the “back-boosting” seen when the vaccine is updated[32] may be due to germinal center formation by memory B cells that are specific to antigenic-sites more conserved between the antigens.

Taken together, our results suggest that children receiving their second dose of the influenza will mount different antibody responses to the vaccine depending on if the vaccine formulation was updated to include antigenically drifted or shifted HA antigens.

## Notes

This original work was supported by the New York Influenza Center of Excellence NIH/NIAID/DMID HHSN272201400005C, the University of Rochester Pulmonary Training Grant T32-HL066988, and the Health Sciences Center for Computational Innovation Pilot Award OP211341.

### Competing Interest Statement

The authors have declared no competing interest.

## REFERENCES

1. Appiah GD, Blanton L, D’Mello T, Kniss K, Smith S, Mustaquim D, et al. Influenza activity – United States, 2014-15 season and composition of the 2015-16 influenza vaccine. MMWR Morb Mortal Wkly Rep. 2015;64: 583–590.

2. Thompson WW, Comanor L, Shay DK. Epidemiology of seasonal influenza: use of surveillance data and statistical models to estimate the burden of disease. Journal of Infectious Diseases. Oxford University Press; 2006;194 Suppl 2: S82–91. doi:10.1086/507558

3. Katz JM, Hancock K, Xu X. Serologic assays for influenza surveillance, diagnosis and vaccine evaluation. Expert Rev Anti Infect Ther. 2011;9: 669–683. doi:10.1586/eri.11.51

4. Trombetta CM, Montomoli E. Influenza immunology evaluation and correlates of protection: a focus on vaccines. Expert Rev Vaccines. 2016;15: 967–976. doi:10.1586/14760584.2016.1164046

5. Fonville JM, Wilks SH, James SL, Fox A, Ventresca M, Aban M, et al. Antibody landscapes after influenza virus infection or vaccination. Science. 2014;346: 996–1000. doi:10.1126/science.1256427

6. Keitel WA, Cate TR, Couch RB, Huggins LL, Hess KR. Efficacy of repeated annual immunization with inactivated influenza virus vaccines over a five year period. Vaccine. 1997;15: 1114–1122.

7. McLean HQ, Thompson MG, Sundaram ME, Meece JK, McClure DL, Friedrich TC, et al. Impact of repeated vaccination on vaccine effectiveness against influenza A(H3N2) and B during 8 seasons. Clin Infect Dis. 2014;59: 1375–1385. doi:10.1093/cid/ciu680

8. Hoskins TW, Davies JR, Smith AJ, Miller CL, Allchin A. Assessment of inactivated influenza-A vaccine after three outbreaks of influenza A at Christ’s Hospital. Lancet. 1979;1: 33–35. doi:10.1016/S0140-6736(79)90468-9

9. Tesini BL, Kanagaiah P, Wang J, Hahn M, Halliley JL, Chaves FA, et al. Broad hemagglutinin-specific memory B cell expansion by seasonal influenza virus infection reflects early-life imprinting and adaptation to the infecting virus. J Virol. 2019. doi:10.1128/JVI.00169-19

10. Gostic KM, Ambrose MR, Worobey M, Lloyd-Smith JO. Potent Protection against H5N1 and H7N9 Influenza via Childhood Hemagglutinin Imprinting. bioRxiv. Cold Spring Harbor Laboratory; 2016;: 061598. doi:10.1101/061598

11. Cobey S, Hensley SE. Immune history and influenza virus susceptibility. Curr Opin Virol. 2017;22: 105–111. doi:10.1016/j.coviro.2016.12.004

12. Smith DJ, Forrest S, Ackley DH, Perelson AS. Variable efficacy of repeated annual influenza vaccination. Proceedings of the National Academy of Sciences. 1999;96: 14001–14006.

13. Li Y, Myers JL, Bostick DL, Sullivan CB, Madara J, Linderman SL, et al. Immune history shapes specificity of pandemic H1N1 influenza antibody responses. J Exp Med. 2013;210: 1493–1500. doi:10.1084/jem.20130212

14. Linderman SL, Chambers BS, Zost SJ, Parkhouse K, Li Y, Herrmann C, et al. Potential antigenic explanation for atypical H1N1 infections among middle-aged adults during the 2013-2014 influenza season. Proc Natl Acad Sci USA. National Academy of Sciences; 2014;111: 15798–15803. doi:10.1073/pnas.1409171111

15. Kelvin AA, Zambon M. Influenza imprinting in childhood and the influence on vaccine response later in life. Euro Surveill. European Centre for Disease Prevention and Control; 2019;24: 722. doi:10.2807/1560-7917.ES.2019.24.48.1900720

16. Gostic KM, Ambrose M, Worobey M, Lloyd-Smith JO. Potent protection against H5N1 and H7N9 influenza via childhood hemagglutinin imprinting. Science. American Association for the Advancement of Science; 2016;354: 722–726. doi:10.1126/science.aag1322

17. Francis T. On the doctrine of original antigenic sin. 1960. doi:10.2307/985534

18. Garten RJ, Davis CT, Russell CA, Shu B, Lindstrom S, Balish A, et al. Antigenic and Genetic Characteristics of Swine-Origin 2009 A(H1N1) Influenza Viruses Circulating in Humans. Science. American Association for the Advancement of Science; 2009;325: 197–201. doi:10.1126/science.1176225

19. Andrews SF, Huang Y, Kaur K, Popova LI, Ho IY, Pauli NT, et al. Immune history profoundly affects broadly protective B cell responses to influenza. Science Translational Medicine. American Association for the Advancement of Science; 2015;7: –316ra192. doi:10.1126/scitranslmed.aad0522

20. Sangster MY, Baer J, Santiago FW, Fitzgerald T, Ilyushina NA, Sundararajan A, et al. B Cell Response and Hemagglutinin Stalk-Reactive Antibody Production in Different Age Cohorts following 2009 H1N1 Influenza Virus Vaccination. Clinical and Vaccine Immunology. 2013;20: 867–876. doi:10.1128/CVI.00735-12

21. Wrammert J, Koutsonanos D, Li G-M, Edupuganti S, Sui J, Morrissey M, et al. Broadly cross-reactive antibodies dominate the human B cell response against 2009 pandemic H1N1 influenza virus infection. J Exp Med. 2011;208: 181–193. doi:10.1084/jem.20101352

22. Nachbagauer R, Choi A, Izikson R, Cox MM, Palese P, Krammer F. Age Dependence and Isotype Specificity of Influenza Virus Hemagglutinin Stalk-Reactive Antibodies in Humans. MBio. American Society for Microbiology; 2016;7: e01996–15. doi:10.1128/mBio.01996-15

23. Ndifon W, Wingreen NS, Levin SA. Differential neutralization efficiency of hemagglutinin epitopes, antibody interference, and the design of influenza vaccines. Proceedings of the National Academy of Sciences. National Acad Sciences; 2009;106: 8701–8706. doi:10.1073/pnas.0903427106

24. Zhang Y, Meyer-Hermann M, George LA, Figge MT, Khan M, Goodall M, et al. Germinal center B cells govern their own fate via antibody feedback. J Exp Med. 2013;210: 457–464. doi:10.1084/jem.20120150

25. Zarnitsyna VI, Lavine J, Ellebedy A, Ahmed R, Antia R. Multi-epitope Models Explain How Pre-existing Antibodies Affect the Generation of Broadly Protective Responses to Influenza. Lauring AS, editor. PLoS Pathog. Public Library of Science; 2016;12: e1005692. doi:10.1371/journal.ppat.1005692

26. Anderson CS, Sangster MY, Yang H, Mariani TJ, Chaudhury S, Topham DJ. Implementing sequence-based antigenic distance calculation into immunological shape space model. BMC Bioinformatics. BioMed Central; 2020;21: 256–13. doi:10.1186/s12859-020-03594-3

27. Anderson CS, McCall PR, Stern HA, Yang H, Topham DJ. Antigenic cartography of H1N1 influenza viruses using sequence-based antigenic distance calculation. BMC Bioinformatics. BioMed Central; 2018;19: 51. doi:10.1186/s12859-018-2042-4

28. Hunt V, Squires RB, Noronha J, Dietrich J, Pickett B, Klem E, et al. Influenza Research Database (IRD): A Web-based Resource for Influenza Virus Data and Analysis [Internet]. Options for the Control of Influenza VII. 2010. pp. 1–1. Available: https://www.fludb.org/brcDocs/posters/Options_poster.pdf

29. Perelson AS, Oster GF. Theoretical studies of clonal selection: Minimal antibody repertoire size and reliability of self-non-self discrimination. Journal of Theoretical Biology. 1979;81: 645–670. doi:10.1016/0022-5193(79)90275-3

30. Chaudhury S, Reifman J, Wallqvist A. Simulation of B cell affinity maturation explains enhanced antibody cross-reactivity induced by the polyvalent malaria vaccine AMA1. J Immunol. American Association of Immunologists; 2014;193: 2073–2086. doi:10.4049/jimmunol.1401054

31. Smith DJ, Forrest S, Ackley DH, Perelson AS. Using lazy evaluation to simulate realistic-size repertoires in models of the immune system. Bull Math Biol. Springer-Verlag; 1998;60: 647–658. doi:10.1006/bulm.1997.0035

32. Fonville JM, Wilks SH, James SL, Fox A, Ventresca M, Aban M, et al. Antibody landscapes after influenza virus infection or vaccination. Science. 2014;346: 996–1000. doi:10.1126/science.1256427

